# Separable neurocomputational mechanisms underlying multisensory learning

**DOI:** 10.1101/2025.11.18.688925

**Authors:** Saurabh Bedi, Ella Casimiro, Gilles de Hollander, Nina Raduner, Fritjof Helmchen, Silvia Brem, Arkady Konovalov, Christian C. Ruff

## Abstract

Efficient control of behavior requires multisensory learning from information distributed across senses. However, most neurocomputational studies have focused on unisensory signals. Here, we identify distinct but interacting neurocomputational mechanisms that support learning of multisensory associations. We designed a task in which behaviorally relevant information was available only from combinations of visual cues with either auditory or tactile cues. In 58 participants undergoing fMRI, we dissociated three processes: multisensory statistical learning (SL), modeled as stimulus-locked Shannon surprise; reinforcement learning (RL), modeled as feedback-locked signed reward prediction errors (RPEs); and feedback-locked unsigned RPEs (uRPEs), reflecting surprise about reward outcomes. Behaviorally, response times scaled with Shannon surprise (SL) while accuracy improved with feedback (RL). Model-based fMRI revealed dissociable but complementary networks: RPEs engaged ventral striatum, vmPFC, and left angular gyrus; surprise recruited bilateral angular gyrus, dlPFC, and precuneus; and uRPEs involved insula, dorsomedial prefrontal, and lateral frontoparietal cortices. Several of these regions extend beyond canonical learning circuits and may contribute to learning when behavior depends on information distributed across sensory modalities. All three networks were modality-general, showing comparable strength for audiovisual and visuotactile learning. Notably, left angular gyrus tracked both Shannon surprise and RPE, suggesting its potential role for integrating structural and value information. These findings indicate that the brain engages distinct but complementary systems for structure-based, reward-based, and outcome-surprise computations. By combining behavioral modeling and fMRI with a novel task design, we provide a framework for dissecting the neurocomputational architecture of multisensory learning.

## Introduction

Most events in daily life engage multiple senses simultaneously: Visual stimuli are paired with sounds, scents, or tactile sensations that the brain seamlessly integrates into a coherent multisensory experience. Reflecting this, neuroscience has increasingly adopted the idea of a multisensory brain [1, 2, 3, 4, 5]. Evidence shows that even brain regions traditionally viewed as unisensory respond to inputs from multiple sensory modalities [6]. Multisensory illusions, such as the ventriloquist and the McGurk effect, further demonstrate that this integration is automatic and can fundamentally alter perception [7]. While the mechanisms of multisensory perception have been extensively studied [8, 9], far less is known about how the brain learns associations, particularly when stimulus- and action-relevant information resides exclusively in the combination of sensory modalities rather than in any individual sense. Understanding the computational and neural principles of multisensory learning remains a gap with implications for many real-world behaviors and cognitive development [10, 11]. For instance, the development of flavor preferences in childhood shapes lifelong eating habits [12] and relies on the integration of smell, taste, and touch, as no single modality suffices to create flavor perception [13, 14]. Language acquisition similarly depends on multisensory learning [11]: reading involves associating visual letters with auditory phonemes and writing involves coordination between visual and tactile information. Impairments in these multi-sensory processes have been associated with developmental dyslexia [15, 16], a learning disability affecting roughly 5 to 10 percent of the population [17]. Together, these observations highlight the need to investigate the computational and neural mechanisms that support learning from multi-sensory information.

Understanding the computational mechanisms underlying multisensory learning begins with identifying the core goals that guide learning more generally. Broadly, learning is shaped by two distinct computational objectives that are each tied to a different source of information. **Reinforcement learning** (**RL**) involves learning to predict rewarding stimuli or actions [18, 19] with the goal of maximizing cumulative reward [20]. In contrast, **statistical learning** (**SL**) refers to the learning of environmental regularities, with the goal of enabling efficient allocation of limited cognitive resources to frequently occurring stimuli [21, 22, 23]. These computations are conceptually distinct: RL is driven by rewards, while SL is driven by the statistical structure of the environment. Here, we examine how each of these learning computations operates in multisensory learning, where both structure and reward are embedded across sensory modalities rather than within unisensory information.

Disentangling RL and SL in multisensory contexts is challenging because these processes often interact. While SL can operate independently of reinforcement, as in classic latent learning [24], it can also support RL by uncovering structure in the environment that facilitates more efficient learning [25, 26, 27, 28, 29]. To examine how these learning computations operate when behavior depends on information distributed across sensory modalities, we designed a novel multisensory task that orthogonalizes RL and SL by separating their informational sources and temporal dynamics. Participants viewed combinations of visual cues with either auditory or tactile signals and judged whether each pairing was correct. Correct responses depended only on the combination of sensory cues rather than on any individual modality. Reinforcement learning was driven by probabilistic feedback, whereas statistical learning was embedded in the frequency with which specific multisensory pairings occurred, providing statistical structure that was irrelevant for reward (see Methods for details). We modeled SL as reward-independent **Shannon surprise** [30] at stimulus onset [31, 32, 33, 34]. Reinforcement learning was modeled using reward prediction errors (RPEs) computed at feedback, capturing the discrepancy between received rewards and expected value estimates derived from the learning model. In addition, we included unsigned RPEs (uRPEs), which index outcome surprise independent of reward valence. Because uRPEs depend on reward feedback like RPEs but quantify statistical surprise like SL, they capture a signal at the intersection of reward-based and structure-based learning. Together, Shannon surprise, signed RPEs, and uRPEs provided complementary measures to dissociate the parallel computations underlying multisensory learning.

This computational and temporal dissociation allowed us to investigate the neural signatures of these learning signals during multisensory learning. Because SL-related surprise occurs at stimulus onset whereas RL-related RPEs and uRPEs occur at feedback, we could temporally separate structure-based and reward-based learning signals in trial-wise fMRI analyses. Previous research in largely unisensory contexts has associated statistical surprise with activity in distributed frontal, posterior parietal regions, and striatal regions [34, 35, 36, 37]. Similarly, signed RPEs are commonly linked to ventral striatal and medial frontal regions, whereas uRPEs have been associated with activity across frontoparietal and insular cortices [38, 39, 37]. However, because these findings are largely derived from unisensory paradigms, and have usually focussed on one type of learning, it remains unknown whether, where, and how the corresponding computations are implemented in parallel when learning depends on associations distributed across multiple senses.

Theoretical accounts of multisensory processing propose several possible architectures for how multisensory computations may be implemented in the brain. One influential framework suggests that certain cortical regions may act as multisensory hubs that integrate information across sen-sory modalities [6]. These hubs may be modality-specific - i.e., engaged differently depending on which specific multisensory combinations are processed - or modality-general, operating similarly across various pairings of different senses. To examine these possibilities, we investigated Shannon surprise, signed RPEs, and uRPEs during multisensory learning across two contexts: audiovisual and visuotactile. Specifically, we asked three key questions: (1) which brain regions track these learning signals during multisensory learning, (2) are these signals modality-specific or modality-general across sensory contexts, and (3) are the different computational components of learning (structure-based (SL), reward-based (RPE), and outcome-surprise–based (uRPE)) supported by shared or distinct neural mechanisms. By combining trial-wise computational modeling with temporally separated stimulus and feedback phases, our design allowed us to isolate each computation and directly test these hypotheses within a unified experimental framework.

## Results

### The experiment

To separate SL and RL during multisensory learning, we developed a two-alternative forced choice task performed by participants (n = 64) across six blocks in an fMRI scanner. In each block, they were presented with one of two multisensory stimulus sets (either audiovisual or visuotactile), each comprising nine unique pairings of three insect images with either three auditory “calls” (beep sequences) or three tactile “dances” (vibration patterns). Participants’ goal was to infer whether each multisensory pairing would attract a mate, mimicking a scientist classifying courtship success in insect species (Fig. 1a, b). Critically, the correct response depended only on the specific multisensory combination, not on any individual unisensory element (Fig. 1c). Feedback was probabilistic: correct pairings typically resulted in successful attraction (+1 reward) but occasionally produced failed attraction (0 reward), whereas incorrect pairings usually resulted in failed attraction but sometimes yielded success (see Methods). This probabilistic feedback ensured that the task remained challenging, promoting continuous RPE signals due to reinforcement learning throughout each block. Participants were compensated with real money based on cumulative rewards they earned during the task.

**Figure 1.**
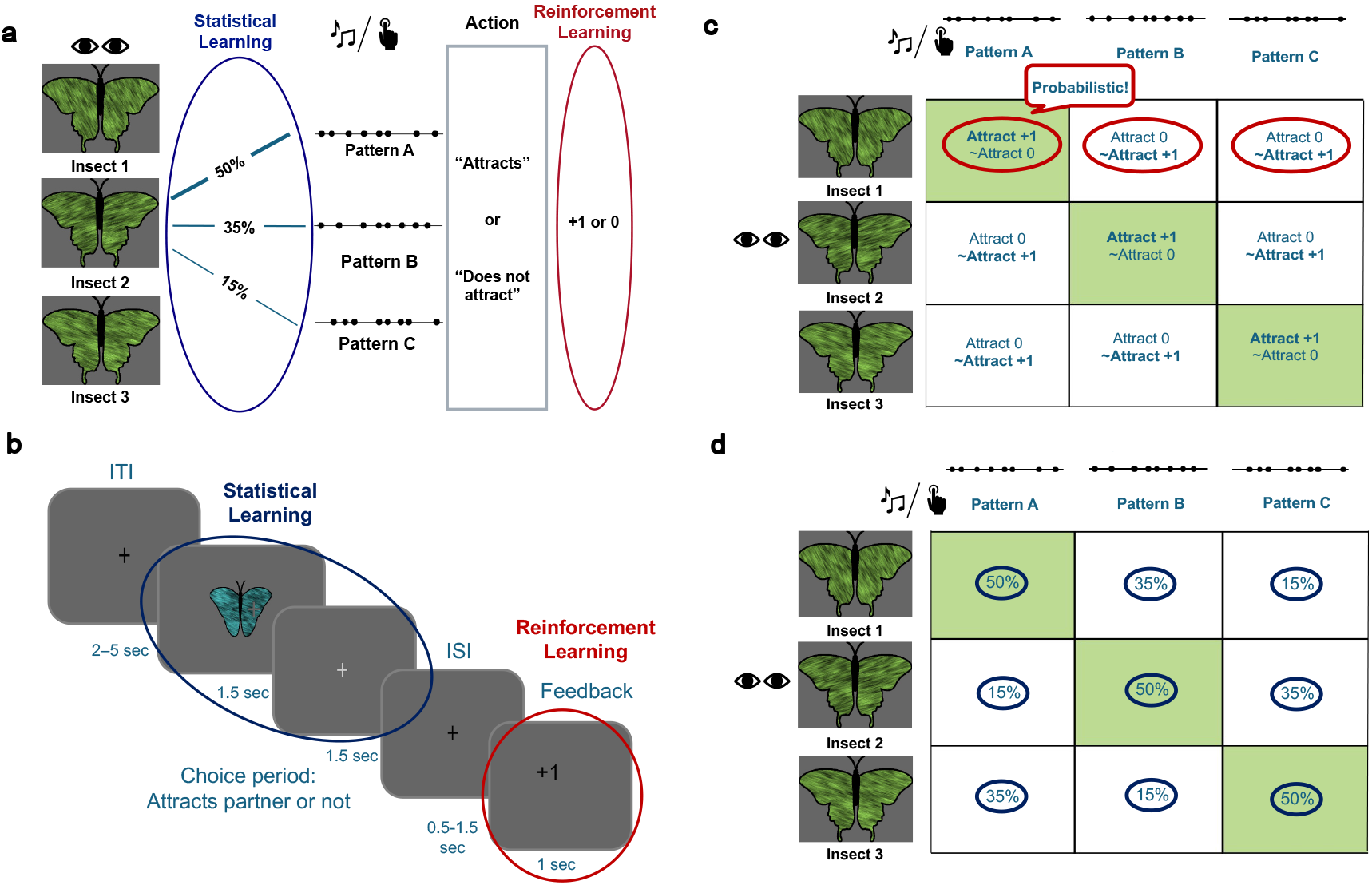
Task design. a, Schematic of the multisensory learning task. Participants acted as “insect scientists” learning which multisensory pairings (image × sound/touch) predicted mating success. b, Trial timing showing separate epochs for stimulus presentation (driving statistical learning) and feedback (driving reinforcement learning). c, Reinforcement structure: probabilistic outcomes (green = “attracts” is rewarded, white = “not attracts” is rewarded) ensured ongoing prediction errors. d, Statistical structure: frequency distribution of multisensory pairings embedding regularities only at the combination level. Together, the task orthogonalized statistical and reinforcement learning signals in both temporal and informational domains.

To isolate SL, we manipulated the frequency with which each multisensory stimulus appeared, embedding statistical structure only in the combinations, not in the visual, auditory, or tactile components individually (Fig. 1d). SL was not required to perform the task or obtain rewards. In contrast, RL was essential: participants had to learn and indicate which combinations led to successful mating. Critically, both successful (green squares) and non-successful (white squares) combinations occurred equally often (50%), meaning that each response (success vs. failure) was correct half the time across all stimuli. This ensured that SL provided no informational advantage for RL. We additionally modeled a third computational signal: the uRPE, which captures the magnitude of outcome surprise regardless of reward valence. While uRPE is computed at feedback alongside signed RPE, it reflects a different computational quantity: the unpredictability of the reward itself. In this sense, uRPE lies at the intersection of RL and SL: like RPE, it is grounded in feedback and value learning, yet like Shannon surprise, it captures statistical unexpectedness, but here in the distribution of reward outcomes. By dissociating these computations in both information content and temporal structure, with SL-relevant surprise occurring at stimulus onset and RL-relevant RPE/uRPE arising after choice (Fig. 1b), our design enabled independent modeling and neural analysis of SL and RL of multisensory stimulus and stimulus-response associations.

## Models of learning

### Reinforcement Learning

We modeled RL to quantify how participants used probabilistic feedback to adapt their (Attract/Don’t attract) choices for each multisensory pairing, and to derive trial-by-trial learning signals (RPE and uRPE) for neural analyses. We used the **Q-learning algorithm** as our base model, in which action values (Q-values) are updated based on the discrepancy between expected value estimates and actual rewards, known as the reward prediction error:

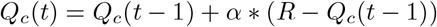

where *Q*_*c*_(*t*) is the Q-value for the chosen action at trial t, R is the received reward (1 or 0), and *α* is the learning rate that determines how strongly RPEs update Q-values. Signed RPEs were computed directly from this update rule, and unsigned RPEs (uRPEs) were defined as the absolute value of the RPE, providing a valence-independent measure of reward-based surprise for neural analysis. It has been shown in purely unisensory contexts that RPEs and uRPEs engage partially distinct neural substrates: while both involve midbrain, striatal, and insular regions, unsigned PEs more consistently activate the cerebellum, dorsolateral and dorsomedial prefrontal cortex, supplementary motor area, supramarginal gyrus, and posterior parietal and temporal cortices, regions implicated in attentional control and salience processing [39, 37]. Including both signals in our analysis therefore allows us to dissociate neural systems involved in value updating from those involved in detecting surprising reward outcomes during multisensory learning.

The **basic Q-learning** model assumes a constant learning rate across all trials [20, 40]. However, participants may have various biases or adopt certain strategies depending on prior expectations or task instructions. To capture these individual differences, we implemented several model variants of Q-learning inspired by the prior literature:

1. Asymmetric Learning Rates (**Asym**): Evidence suggests that participants may learn differently from positive and negative RPEs [41, 42, 43, 44]. To capture this asymmetric learning, we implemented models with distinct learning rates for positive and negative RPEs.
2. Transfer Learning Models (**Transfer**): The task instructions explicitly stated that each insect had only one correct call or dance. This means that if a participant received a reward for pairing insect A with call 1 (A1), they could infer that other pairings involving the same insect (e.g., A2, A3) or the same call (e.g., B1, C1) are likely incorrect. This rule-based structure, which was available from the instructions but not through direct feedback, creates opportunities for inference beyond the presented trial. We modeled this as a structured “transfer” mechanism that allowed RPEs to update not only the chosen option but also unchosen but logically related options [45].
3. Initial Bias Model (**Vinit**): Participants might begin the task with preexisting biases toward one response option (e.g., left/right). We modeled this by assigning different initial values to actions, allowing for baseline preferences that could influence early behavior.

We also fit combinations of these models (e.g., Transfer+V_init_), yielding eight candidate models in total (see Methods). Of these, five showed acceptable parameter recovery (*r >* 0.6) and were used for final model selection. Across participants, 31 were best fit by the basic Q-learning model, 16 by the Transfer model, and 11 by the Asymmetric model, while neither the Vinit model nor the Transfer+V_init_ combination best fit any participant (see Methods for details). Each participant’s best-fitting model per block was then used to compute trial-by-trial estimates of RPE and uRPE for neural analyses.

Action selection on each trial was modeled using a softmax function:

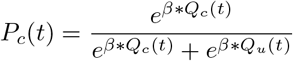

where *Q*_*c*_ is the value of the chosen response and *Q*_*u*_ is the value of the unchosen response. The inverse temperature parameter *β* scales the level of randomness in choices and was fit for each individual for each block.

### Statistical Learning

We modeled SL to test whether participants implicitly tracked the frequency structure of multisensory combinations, despite its irrelevance for reward, and to obtain a stimulus-locked, trial-by-trial index of structure-driven surprise for behavioral and fMRI analyses. We used a simple Bayesian observer that maintains beliefs over the nine combinations (uniform prior at block start) and updates them from observed frequencies [31, 32, 33, 34]:

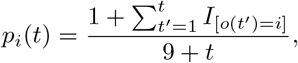

where *p*_*i*_(*t*) is the belief that stimulus *i* occurs on trial *t* and 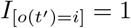 if *i* was observed on trial *t*^*′*^. The SL regressor is the Shannon surprise at stimulus onset,

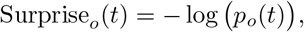

which quantifies how unexpected the presented combination *o(t)* is given current beliefs. Prior work has shown that such signals can modulate behavioral response times and neural responses, even in the absence of explicit rewards [34]. Because regularities were confined to joint combinations and were balanced across actions and modalities, this stimulus-locked signal is orthogonal to RL. In contrast to uRPE, which captures surprise about reward outcomes at feedback, Shannon surprise quantifies surprise about stimulus states at onset.

### Behavioral results

Our behavioral data indicate that participants were engaged in both reinforcement and statistical learning during the task. As shown in Fig. 2a, accuracy increased steadily over trials in both audiovisual and visuotactile blocks, reflecting reinforcement learning from the reward feedback. The probabilistic reward structure encouraged continued exploration and prevented early learning saturation. To test the relationship between surprise and response times, we fit a mixed-effects regression model that included surprise, Q-values, and subject-level random intercepts as predictors. Surprise was positively associated with response time in both modalities (audiovisual: *β* = 0.023, one-sided p = 0.013; visuotactile: *β* = 0.017, one-sided p = 0.037). This effect, visualized in Fig. 2b, shows that responses were slower for less frequent stimulus combinations, suggesting that participants were sensitive to the statistical structure of the stimulus distribution.

**Figure 2.**
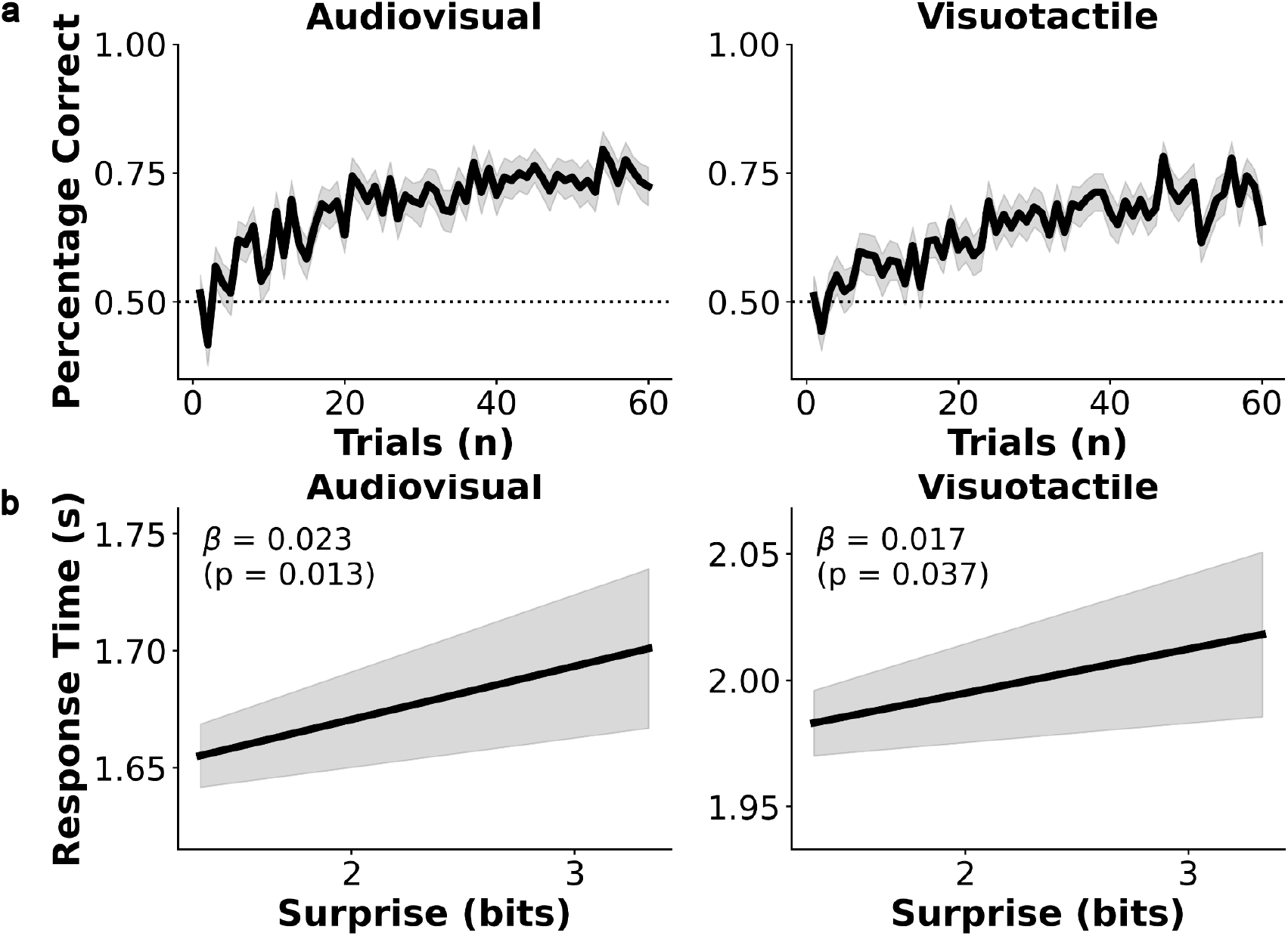
Behavioral signatures of reinforcement and statistical learning. a, Group-averaged learning curves showing accuracy across trials (smoothed, ±SEM) for audiovisual and visuotactile conditions. Increasing accuracy indicates reinforcement learning driven by feedback-based reward prediction errors. b, Response times as a function of Shannon surprise (computed at stimulus onset) showing the hypothesized longer latencies for rare combinations, consistent with implicit statistical learning. Each line represents the fixed-effect slope from a mixed-effects model controlling for subject-level random intercepts. Together, these effects confirm that participants engaged both feedback-based and structure-based learning mechanisms.

### Neural underpinnings of the two types of multisensory learning

Having established distinct behavioral and computational signatures of RL and SL, we next examined whether these two learning processes are implemented in separate neural systems. Specifically, we tested three key questions: (1) whether multisensory SL and RL signals are encoded in distinct brain regions, (2) whether their neural substrates are modality-general or modality-specific, and (3) whether any identified brain areas extend beyond regions commonly reported in unisensory learning studies. To address these questions, we conducted a whole-brain model-based fMRI analysis [46], using trial-wise RPEs, uRPEs and Shannon surprise as parametric modulators for RL and SL. Standard nuisance regressors (motion, physiological noise) were also included in the general linear model, and all results were thresholded using non-parametric cluster-based inference (SnPM13).

In our first whole-brain GLM analysis, we collapsed across audiovisual and visuotactile conditions to target domain-general learning signals (domain-specific signals are examined in a second set of analyses). These first analyses revealed that the three learning signals (RPE, Shannon surprise, and uRPE) were each associated with largely non-overlapping brain networks, supporting the idea that distinct neural architectures support reinforcement and statistical learning. Trial-by-trial RPEs during the feedback phase were correlated with activity in the ventromedial prefrontal cortex (vmPFC) and ventral striatum (caudate nucleus, putamen), all regions consistently implicated in unisensory reinforcement learning, particularly in value representation and reward prediction error encoding across modalities [47, 48, 49, 50, 51, 52]. Additional RPE-related activation appeared in visual association areas (fusiform, lingual, and middle occipital gyri; see Fig. 3 and Table 1), likely reflecting visual stimulus-dependent value modulation or attentional processing [53, 37, 38]. Notably, activity in left angular gyrus also tracked RPEs, hinting at a potential region beyond classical reward circuitry. This region has so far only been reported to be involved in static reward anticipation [54, 55], but not in representation of dynamically evolving RPEs [37, 53, 38]. Inspection of the reported coordinates and available activation maps from these meta-analyses of unisensory RPE processing revealed no spatial overlap with the present activation cluster, sug-gesting that its involvement may reflect computations that are relevant when learning depends on information distributed across modalities or cues.

**Figure 3.**
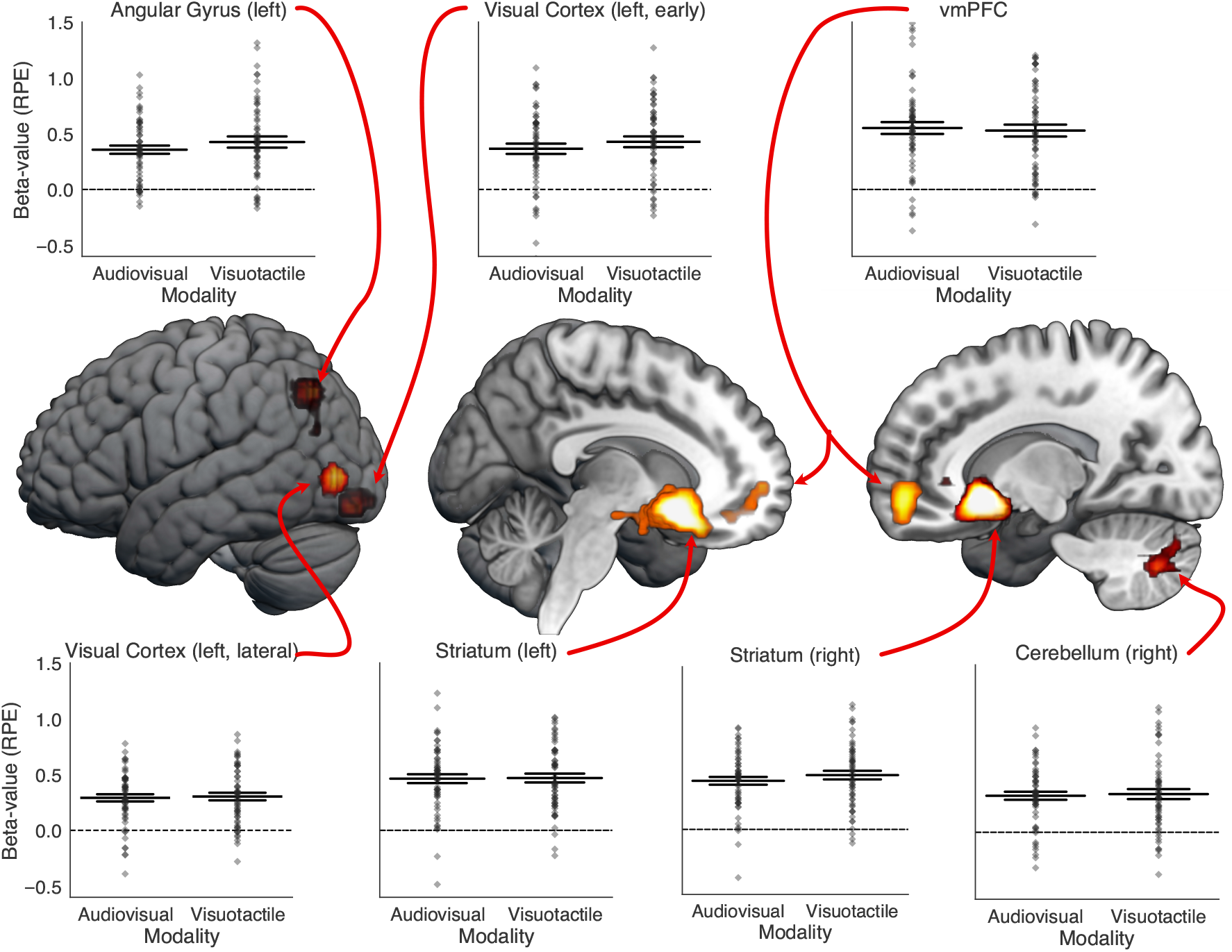
Neural correlates of reward prediction errors (RPEs). The brain images in the center display regions where signed RPEs, modeled as a parametric modulator at feedback onset, positively correlated with BOLD activation. RPE reflects the signed difference between expected and received feedback (reward-dependent) indexing reinforcement learning. Statistical maps were thresholded at a cluster-forming *t >* 8.0 and inference was performed at cluster-level *p <* .05, FWE-corrected using SnPM, and rendered on the MNI152 standard brain. Significant activations were observed in the bilateral striatum, ventromedial prefrontal cortex (vmPFC), early lateral left visual cortex, left angular gyrus, and right cerebellum. The accompanying strip plots display the mean beta estimates of the RPE contrast across all voxels within the thresholded cluster in MNI space, with each diamond marker representing data from an individual participant. Estimates are shown separately for audiovisual and visuotactile learning, revealing that RPE-related responses did not differ significantly between different combinations of sensory modalities. The widest horizontal line indicates the group mean, whereas the more narrow lines indicate a distance of one standard error of the mean (SEM).

In line with the assumption that separate neural systems mediate RL vs SL, Shannon surprise during stimulus presentation engaged a separate set of regions, including bilateral angular gyri, the left precuneus, and the left dorsolateral prefrontal cortex (dlPFC; see Fig. 4 and Table 2). To our knowledge, only a handful of studies have examined the neural correlates of Shannon surprise in unisensory contexts [37, 36, 35, 34] and no large-scale meta-analyses are available. Despite this limited evidence base, our findings partially converge with previous reports: all four previous studies also identified a parietal cluster near to (or extending into) the angular gyrus, and two of the four studies also linked the dlPFC to the Shannon surprise, albeit in the right hemisphere [37, 34]. Notably, none of these studies reported activation in the precuneus area identified here, suggesting that this region may contribute to a modality-independent representation of multisensory surprise. In any case, the absence of spatial overlap with RPE-related regions further supports the conclusion that statistical and reinforcement learning rely on distinct neurocomputational pathways.

**Figure 4.**
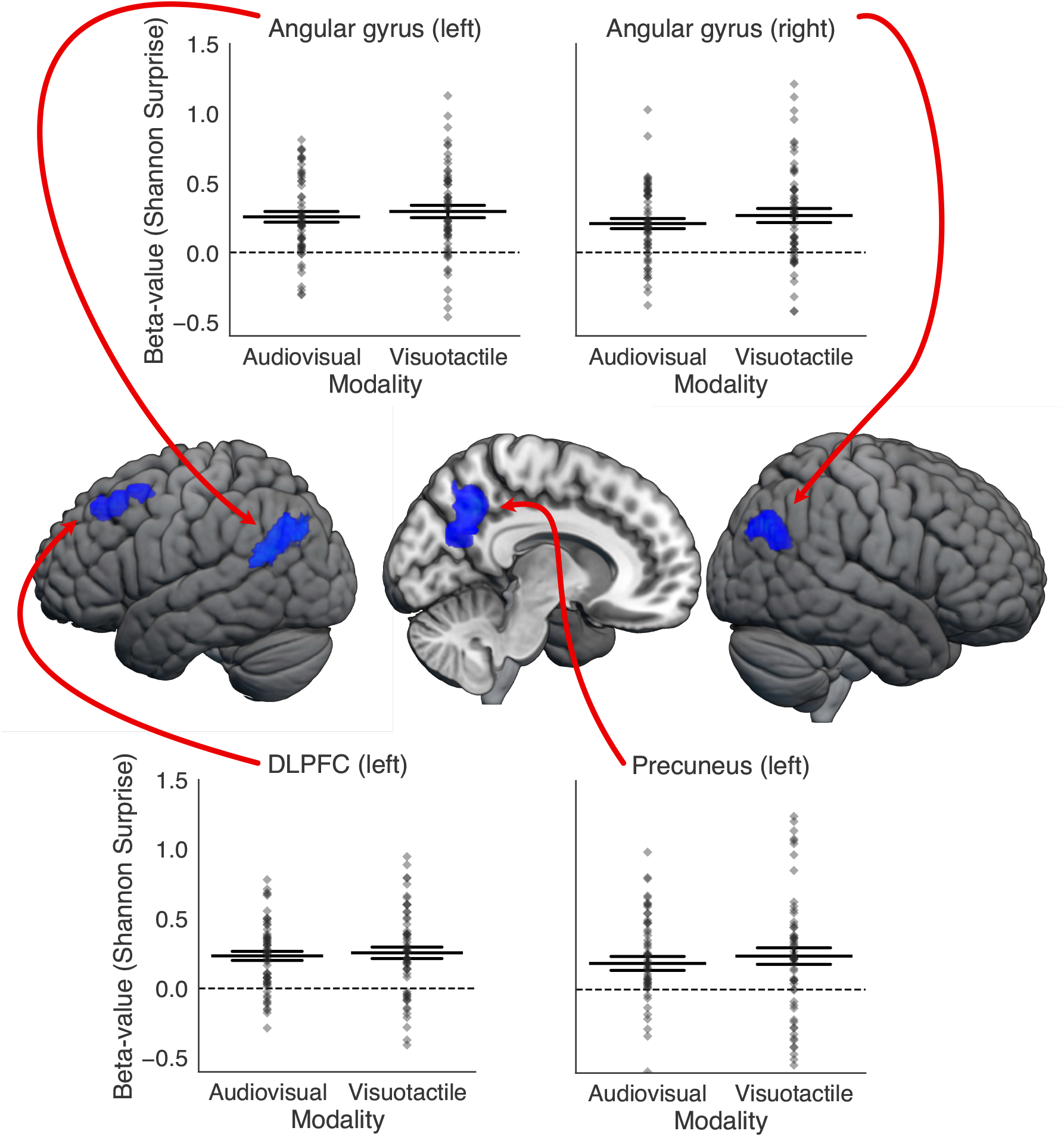
Neural correlates of surprise. The brain images in the center display regions where the Shannon surprise, modeled as a parametric modulator at stimulus onset, positively correlated with BOLD activation. The Shannon surprise quantifies the unexpectedness of the stimulus and indexes statistical learning. Statistical maps were thresholded at a cluster-forming *t >* 3.1 and inference was performed at *p <* .05, FWE-corrected using SnPM, and rendered on the MNI152 standard brain. Surprise significantly correlated with brain activation in the bilateral angular gyrus, left dorso-lateral prefrontal cortex (dlPFC), and left precuneus. The accompanying strip plots display the mean beta estimates of the RPE contrast across all voxels within the thresholded cluster in MNI space, with each diamond marker representing data from an individual participant. Estimates are shown separately for audiovisual and visuotactile learning, revealing that RPE-related responses did not differ significantly between modalities. The widest horizontal line indicates the group mean, whereas the more narrow lines indicate a distance of one standard error of the mean (SEM).

To further test the dissociability of learning signals, we modeled uRPE, a valence-independent measure of outcome surprise computed at feedback. Conceptually, uRPE bridges RL and SL: like RPE, it depends on rewards, and like Shannon surprise, it reflects deviation from reward expectation. Given this conceptual overlap, we examined whether uRPE would recruit a distinct or shared network relative to RPE and surprise signals. In line with the former option, neural responses to uRPEs revealed a third, largely distinct network, with significant activation in the right dorsomedial PFC, bilateral insula, and a lateral frontoparietal circuit including inferior parietal lobule, middle frontal gyrus, orbital inferior frontal gyrus, and superior frontal sulcus (see Fig 5 and Table 3). This network aligns in part with prior reports of uRPE-related activation in the insula, dmPFC, and inferior parietal regions [37, 39], supporting their role in tracking reward salience and behavioral relevance. However, compared to any previous findings, the frontal clusters identified in our study appear to be more lateral and ventral than any previously observed activations. This spatial shift may reflect the additional computational demands of our multisensory task, in which feedback integration and prediction updating occur across, rather than within, sensory modalities [6]. In such contexts, the brain must not only evaluate prediction errors but also determine which sensory stream contributed to the unexpected outcome and how these streams jointly inform future expectations. The recruitment of lateral and ventral frontal regions could therefore indicate enhanced engagement of cognitive control [56] and multisensory integration processes [57] needed to arbitrate between converging sensory inputs during outcome evaluation. Together, these results support the idea that uRPE engages a third, largely dissociable network, which is functionally distinct from both RPE- and surprise-related circuits, even when operating within the same multisensory learning task.

**Figure 5.**
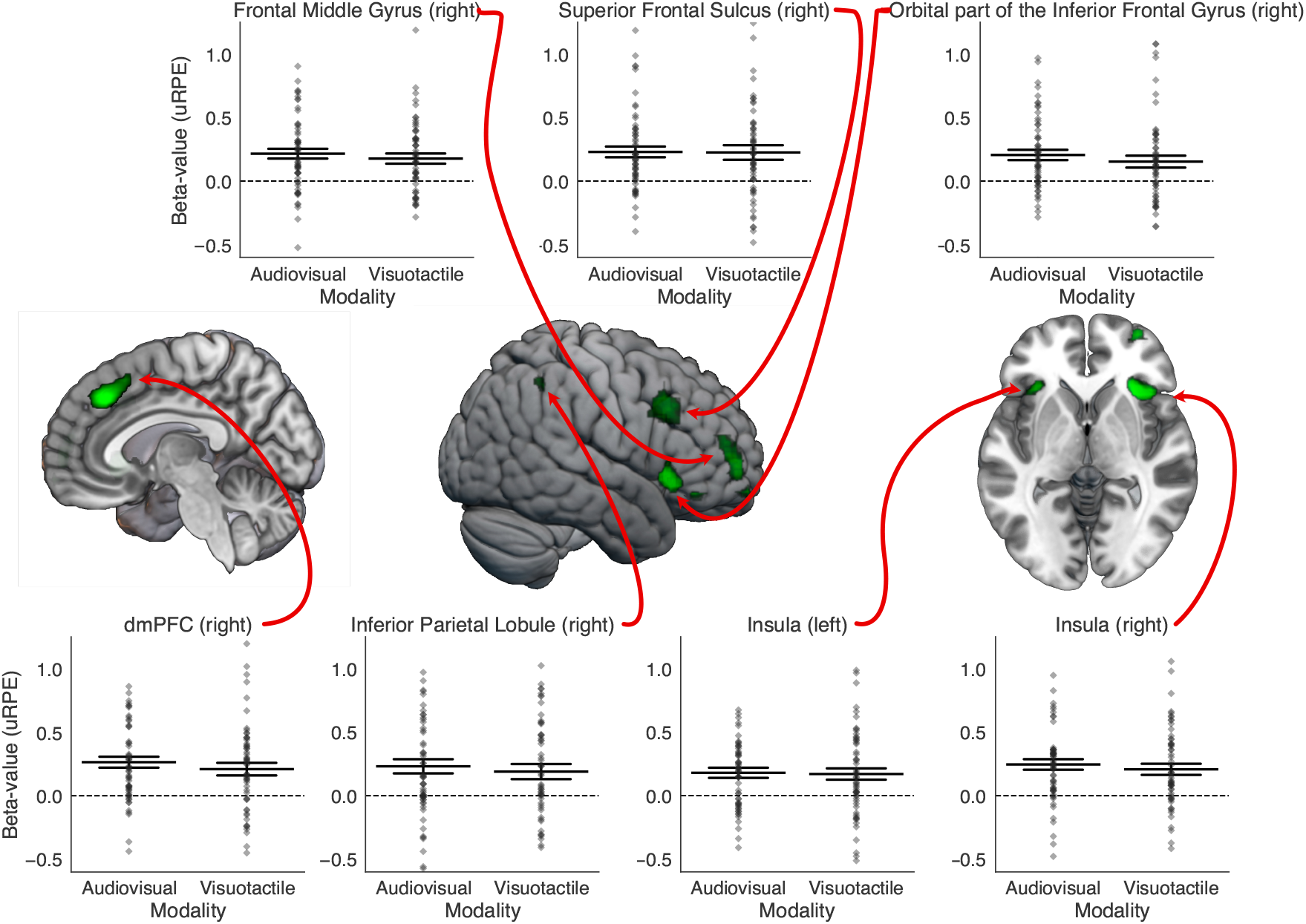
Neural correlates of unsigned reward prediction errors (uRPEs). The brain images in the center display regions where unsigned RPEs, modeled as a parametric modulator at feedback onset, positively correlated with BOLD activation. uRPEs capture the magnitude of outcome surprise regardless of feedback valence. Statistical maps were thresholded at *t >* 4.0 and then *p <* .05, FWE-corrected using SnPM, and rendered on the MNI152 standard brain. Significant activations were observed in the right inferior, middle and superior frontal cortex, right dorso-medial prefrontal cortex (dmPFC), right inferior parietal lobe, and bilateral insula. The accompanying strip plots display the mean beta estimates of the RPE contrast across all voxels within the thresholded cluster in MNI space, with each diamond marker representing data from an individual participant. Estimates are shown separately for audiovisual and visuotactile learning, revealing that RPE-related responses did not differ significantly between modalities. The widest horizontal line indicates the group mean, whereas the more narrow lines indicate a distance of one standard error of the mean (SEM).

In a second set of whole-brain GLM analyses, we tested whether RL and SL are supported by modality-general or modality-specific mechanisms. We computed whole-brain contrasts for each learning signal (RPE, Shannon surprise, and uRPE), comparing visuotactile and audiovisual conditions (visuotactile *>* audiovisual and vice versa). No significant clusters emerged for any of the three learning signals. This suggests that, at the functional and anatomical resolution of our data, there is no evidence for modality-specific coding, neither in terms of increased or decreased activity in the identified regions, nor in the spatial distribution of the regions coding for these signals. Given the absence of significant modality-specific effects in the whole-brain analysis, we proceeded to examine whether the magnitude of learning signal coding in the already-identified domain-general learning ROIs differed across modalities. To this end, we extracted beta weights from the significant ROIs identified in the main contrasts (i.e., all voxels within clusters surviving the cluster-forming threshold). These beta weights showed comparable activation levels across both multisensory modalities (see swarm plots in Fig. 3, Fig. 4, and Fig. 5). In sum, these results suggest that the brain uses shared, domain-general systems to encode RPEs, surprise, and outcome uncertainty for multisensory learning, even when the precise sensory input channels differ, highlighting the generality and contextual flexibility of learning circuits in multisensory environments.

Finally, we explored whether any of the regions we observed may be involved in processes important for multisensory learning, beyond their role in general RL and SL (Fig. 6). While most brain regions revealed in our GLM analyses overlap with known SL and RL networks [39, 53, 38, 34, 35, 36, 37], several regions are not typically observed to be involved in learning in unisensory contexts (indicated by dotted outlines in Fig. 6). Most notably, despite the largely distinct regions underlying RPE and Shannon surprise, the **left angular gyrus** consistently tracked both these signals. Prior studies have reported neighboring parietal regions like the inferior and superior parietal lobules for surprise [35, 34, 36], but have not implicated the angular gyrus as a locus for statistical learning [37, 39, 53, 38]. Similarly, while the inferior parietal lobule has been associated with RPEs in an extensive meta-analysis [37], the clusters from our findings and the meta-analysis do not overlap. In contrast, our findings show robust angular gyrus engagement in both statistical and reinforcement-based multisensory learning, suggesting that this region, sitting at the intersection of the higher order visual, auditory, and tactile cortices [58], may play a role in integrating structural and value-related information during multisensory learning.

**Figure 6.**
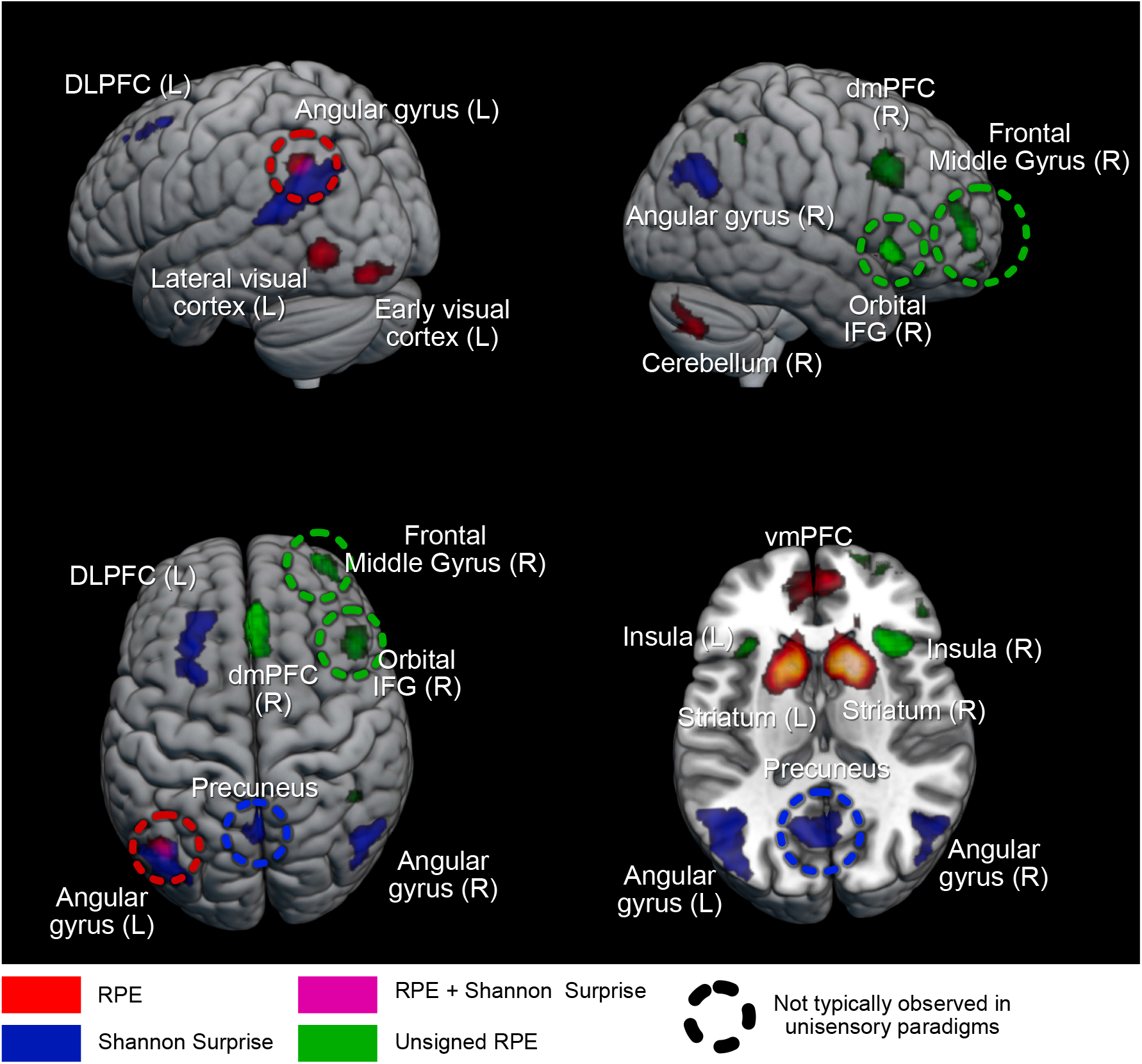
Dissociable neural networks underlying multisensory learning. Distinct brain regions encode reward prediction errors (RPE, red), Shannon surprise (blue), and unsigned reward prediction errors (uRPE, light green) during multisensory learning. The three networks show minimal overlap, highlighting specialized yet complementary systems for reinforcement learning, statistical learning, and outcome surprise. Regions outlined with a dotted line are not typically reported in unisensory learning tasks, suggesting involvement in computations that are relevant to learning multisensory associations. Brain maps are rendered on the MNI152 standard brain template.

We also observed surprise-related activity in the precuneus, a region widely recognized as a major cortical hub that integrates information from multiple sensory and cognitive networks [59, 60,, despite the precuneus not reported in prior unisensory SL studies [35, 34, 36, 37]. This suggests that the precuneus may also provide a modality-independent estimate of multisensory surprise. Regarding uRPE, several regions we observed (bilateral insula, inferior parietal lobule, superior frontal sulcus, and dmPFC) have been linked to uRPEs before [37, 39]. However, the clusters in the frontal middle gyrus and orbital inferior frontal gyrus do not overlap with regions described in previous literature. Furthermore, regions typically associated with uRPEs in unisensory contexts - such as the striatum, cingulate cortex, and temporal regions - were not found to be reliably engaged in our study. Our results thus reveal distinct networks that underlie the three types of multisensory learning; each of these comprise areas typically implicated in unisensory learning but crucially also other areas that appear to be functionally relevant when learning depends on multisensory associations.

## Discussion

Everyday learning often depends on non-redundant information distributed across multiple senses (e.g., the integration of taste and smell in perceiving flavor), yet most work has characterized learning with unisensory stimuli. Existing studies have modeled reinforcement learning with signed reward prediction errors [40, 62], which drive learning behavior and neural activity in structures like ventral striatum/vmPFC [50, 47, 51, 52, 63]. In parallel, stimulus-structure-driven SL is often modeled as Shannon surprise and has been shown to drive both behavior and cortical responses in largely unisensory paradigms [64, 34, 35, 36]. To extend these approaches to situations in which informative structure is distributed across multiple senses, we designed a task in which learning-relevant information existed only in multisensory stimulus combinations rather than in any single modality. This design enabled temporal and informational dissociation between SL - indexed by stimulus-locked Shannon surprise - and RL indexed by feedback-locked RPE. In addition, participants’ evolving expectations about reward outcomes generated a third signal: uRPEs, which index outcome surprise and lie at the computational intersection of RL and SL. uRPEs share RL’s dependence on expected rewards and SL’s emphasis on deviation from these expectations, and are also known to modulate both learning behavior and neural responses in unisensory paradigms [65, 39, 37]. By isolating stimulus-driven structure, outcome-driven learning, and performance-contingent outcome surprise, our task and model-based fMRI results [46] thus provide a principled framework for studying how the brain supports multiple forms of multisensory learning within a unified context.

Importantly, the present findings should not be interpreted as evidence that the neural systems we identified are specific to multisensory learning per se. Rather, our task isolates learning signals in a situation where behaviorally relevant information is distributed across multiple cues, such that no single sensory channel alone can guide behavior. Multisensory environments provide a natural instance of this problem, but similar computational demands may arise in all situations where informative structure is distributed across different stimulus features or dimensions. In this sense, multisensory learning provides a tractable model system for studying how the brain integrates structural and value-based information across multiple sources.

Behaviorally, we observed the expected signatures of SL and RL. Response times increased with trial-wise Shannon surprise (hallmark of SL), indicating higher response times for rarer multisensory stimuli, and accuracy improved with reward feedback [34, 51]. Neurally, model-based fMRI revealed largely non-overlapping systems for each signal. RPEs recruited classical reward-related regions (ventral striatum, vmPFC) [47, 52, 63, 37, 38, 39], while Shannon surprise engaged a separate network in parietal and frontal cortices [36, 35]. uRPEs were encoded in a third, dissociable network involving the insula, dorsomedial prefrontal cortex, and frontoparietal regions [39, 37]. These brain regions overlap with both the salience network and the frontoparietal network, indicating that uRPEs may engage systems responsible for detecting behaviorally relevant events and implementing adaptive cognitive control [66, 67]. This suggests that the brain not only learns from environmental structure and feedback, but also monitors surprising feedback to potentially guide attentional shifts or cognitive control. Despite their distinct circuitry, all three signals (RPE, Shannon surprise, and uRPE) were encoded in modality-general networks, with no differences across audiovisual and visuotactile conditions. These findings suggest that the brain deploys specialized but flexible learning systems to support complex learning from multisensory input across various multisensory contexts. Nonetheless, modality-specific anatomical organization may exist at a finer scale than resolved by our standard-space fMRI analysis and should be addressed by future work using high-resolution methods [68, 69].

Among the brain regions engaged by our task, the left angular gyrus was notable in tracking both RPE and Shannon surprise, despite the otherwise clear dissociation between multisensory SL and RL signals at the whole-brain level. This pattern suggests that this region contributes to integrating structure- and value-based learning signals when information is distributed across modalities. While neighboring parietal areas have previously been linked to either SL or RL in unisensory contexts [35, 36], the angular gyrus itself is not typically highlighted in meta-analyses of either computation. Its involvement here is consistent with proposals that angular gyrus functions as a multimodal integration hub [58]. Anatomically, the region is densely connected with higher-order visual, auditory, and somatosensory association cortices [58, 70, 71], positioning it well to integrate cross-modal information. In large-scale network terms, the angular gyrus forms part of the salience and frontoparietal control networks, which are typically engaged in detecting and reorienting to behaviorally relevant sensory events [58]. These converging properties make it a plausible cortical locus for externally driven multisensory learning and integration. The angular gyrus has also been implicated in higher-level semantic processing [72, 73], and altered activation in temporoparietal regions surrounding the angular gyrus has been reported in reading and language disorders [74, 75, 76, 77]. However, meta-analyses did not link angular gyrus activation specifically to dyslexia [78, 79]. While the regions identified here may therefore not correspond directly to the core neural abnormalities underlying dyslexia, the angular gyrus likely contributes to integrating perceptual inputs—such as written or spoken words—with higher-level semantic representations, a process critical for reading acquisition. Future studies combining the present paradigm with causal methods such as transcranial magnetic stimulation (TMS) could directly test this possibility.

We also observed surprise-related activity in the **precuneus**, a region densely interconnected with parietal, temporal, and prefrontal association cortices and often proposed to be a cortical hub that integrates information from multiple sensory and cognitive networks [60, 61, 59]. Interestingly, this region has not yet been linked to Shannon surprise [35, 34, 36, 37]. The presence of surprise-related signals in the precuneus may reflect the updating of internal, higher-order models that link multisensory inputs with self-referential or contextual expectations according to predictive coding frameworks [80]. In large-scale network terms, the precuneus is a core node of the default mode network, which supports internally oriented inference, mental model updating, and self-referential evaluation [59, 60, 81]. Its recruitment therefore complements the engagement of the angular gyrus, which has been more closely associated with salience and frontoparietal control networks supporting externally directed attention. In parallel, uRPE-related activation in frontal regions has not been consistently linked to feedback-driven surprise, unlike the lateral fronto-parietal cortex more commonly associated with the computation [39, 37].

Together, our findings suggest that multisensory learning may recruit neural circuits that extend beyond those typically reported in unisensory learning contexts. Notably, atypical activation patterns in the regions identified here have also been reported in developmental disorders affecting learning and cognitive control. In both Attention Deficit Hyperactivity Disorder (ADHD) [82, 83] and Autism Spectrum Disorder (ASD) [84], hypoactivation in frontal regions has been linked to deficits in executive functioning and attentional control. In dyslexia, frontal activation patterns are more heterogeneous, with both hypo- and hyperactivation reported across studies [78, 85]. The observed hypoactivation has been interpreted as reflecting executive dysfunction [86] and difficulties in accessing lexical and sublexical phonological representations [78], and it has also been associated with the high comorbidity between dyslexia and ADHD [86]. Conversely, hyperactivation has been suggested to reflect articulatory compensatory mechanisms [78] or a broader spatial distribution of the reading network [87].

Of particular relevance, the inferior frontal gyrus, together with the posterior superior temporal sulcus, occipito-temporal cortex, and inferior parietal lobe, forms the core network underlying audiovisual letter–speech sound learning and integration [88]. Both children and adults with dyslexia exhibit reduced activation within this network, as well as impaired functional and structural connectivity among its constituent regions [88]. Atypical hyperactivation of the precuneus has likewise been observed in ASD [84, 89] and ADHD [89], likely reflecting insufficient deactivation of the default mode network. In dyslexia, by contrast, reduced precuneus activation relative to typical readers has been reported [85, 90], suggesting increased cognitive effort and compensatory recruitment during reading-related tasks [90, 79]. Although the present study did not directly investigate these disorders, our findings highlight that neural systems supporting learning from distributed sensory information overlap with circuits known to be affected in atypical development.

Several caveats warrant mention. First, as with all fMRI studies, the present findings are correlational and cannot establish causal involvement of the identified regions. Future work using causal approaches such as TMS [91, 92, 93] or lesion analyses [94] will be required to test whether these regions are necessary for multisensory learning. While superficial cortical areas such as the angular gyrus are accessible to TMS, deeper or midline structures may require emerging techniques such as focused ultrasound [95, 96] or deep-TMS [97]. Studying developmental and aging populations may also reveal how multisensory learning and the integration of value and structure evolve across the lifespan [98]. Finally, extending this framework to other ecologically relevant domains, such as speech perception, reading, or sensorimotor coordination, may clarify how the multisensory learning computations identified here extend to other cognitive functions. Applying the present modeling approach in these contexts could help determine whether the same neural computations that integrate structure and feedback across modalities also support language and reading acquisition, and whether they are altered in atypical development. In any case, our experimental approach and findings provide a starting point for future studies on how the brain integrates structural and value-based information when learning associations distributed across multiple sensory sources.

## Acknowledgements

We are grateful to C. Schnyder, K. Treiber, and M. Moisa at the Zurich Center for Neuroeconomics for their excellent support with recruitment and participant coordination. We thank the staff of the Laboratory for Social and Neural Systems Research for technical assistance throughout data acquisition.

This research was supported by the University Research Priority Program (URPP) Adaptive Brain Circuits in Development and Learning (AdaBD) at the University of Zurich. C.C.R. was additionally supported by the Swiss National Science Foundation (SNSF; grant no. 100019L-173248).

## Methods

### Participants

Sixty-four right-handed participants (21 women; age range 18–30 years, mean age = 23.3) took part in the fMRI study. All participants were screened for MRI compatibility prior to inclusion and reported no psychiatric or neurological disorders or need for visual correction. Participants were students at the University of Zurich or ETH Zurich, with those enrolled in Economics, Psychology, or Computer Science excluded from participation. Six participants were removed from the analyses due to scanner-related problems or excessive head motion, resulting in a final sample of 58. All procedures conformed to the Declaration of Helsinki and were approved by the Ethics Committee of the Canton of Zurich.

### Procedure

The fMRI study was conducted at the Laboratory for Social and Neural Systems Research (SNS Lab) of the University Hospital Zurich. Each session lasted no more than 2 hours and 15 minutes. Upon arrival, participants completed MRI safety screening and provided written informed consent. They then proceeded to a behavioral testing room, where they read instructions for the multisensory learning task, answered comprehension questions, and received additional information on MRI safety. Participants subsequently performed the multisensory learning task inside the MRI scanner (Philips ACHIEVA 3T), during which both behavioral and neural measures were collected. Visual stimuli were projected onto a screen and viewed via a mirror mounted on the head coil. Auditory stimuli were delivered using MRI-compatible headphones (MR Confon GmbH, www.mr-confon.de), and tactile stimuli were applied to the left index finger using a piezoelectric stimulator (mini-PTS, Dancer Design, http://www.dancerdesign.co.uk/). Eye movements were recorded with an MR-compatible infrared Eyelink II CL v.4.51 eye-tracker (SR Research Ltd), and physiological data were obtained using a breathing belt and an MR-compatible ECG device. In addition to functional imaging, we acquired a high-resolution anatomical scan and diffusion tensor imaging (DTI) data. Participants received a base payment of 50 CHF, plus a performance-dependent bonus of up to 48 CHF.

### Multisensory learning task

Participants were instructed to play a game in which they took the role of a scientist studying newly discovered insects. On each trial, a visual image of an insect was paired with either an auditory “call” (a sequence of beeps presented over headphones) or a tactile “dance” (a sequence of vibrations delivered to the left index finger). Participants judged whether the multisensory pairing would attract a mate (“attract” vs. “no attract”) with a button press.

#### Visual stimuli

Each block featured three insect “species,” distinguished solely by the orientation of a textured noise pattern superimposed on a butterfly-shaped silhouette (Fig. 1a). The textures were generated by filtering pink noise with von Mises orientation filters, yielding three distinct orientation-defined patterns at approximately 22.5^°^, 67.5^°^, and 112.5^°^. These angles were chosen to be equally spaced away from the cardinal axes. Images were exported at high resolution (1200 dpi), presented centrally on a gray background. Importantly, species identity could be determined from this single visual feature (orientation).

#### Auditory and tactile stimuli

Auditory “calls” and tactile “dances” were constructed as sequences of discrete bursts presented within a fixed 1.5 s stimulus window. Each pattern contained 9 bursts: 8 tone (or vibration) bursts separated by variable inter-burst intervals, plus a final burst, such that the entire sequence always filled 1.5 s. Each burst lasted 15 ms. For auditory stimuli, bursts were sine tones (delivered via MRI-compatible headphones), while for tactile stimuli, bursts were square pulses delivered to the left index finger via a piezoelectric stimulator (mini-PTS, Dancer Design; 48 kHz sampling rate). For both modalities, three distinct temporal patterns were created per set by varying the distribution of inter-burst intervals while keeping total duration fixed. This ensured that tactile and auditory stimuli were structurally analogous and directly comparable across modalities.

#### Task structure and timing

The experiment consisted of six blocks (runs), alternating between audiovisual (calls) and visuotactile (dances) modalities (3 blocks each). The starting modality was randomized across participants. Each block contained 60 trials. Trial timing was as follows: the multisensory stimulus was presented for 1.5 s (during which responses were recorded), followed by a jittered inter-stimulus interval (0.5–1.5 s), then feedback displayed for 1 s. Inter-trial intervals were jittered between 3–5 s to optimize design efficiency and minimize correlations between parametric regressors. A response-time window of up to 3.0 s was enforced (responses after this window were treated as missing). The total task duration was approximately 54 minutes.

### Reinforcement-learning (feedback) contingencies and calibration

Each insect species had exactly one “high-success” pairing (i.e., one specific call or dance most likely to attract a mate), whereas the other two pairings for that species had much lower success rates. Feedback was probabilistic: correct pairings led to “+1” (green tick) most of the time, while incorrect pairings yielded “0” (red cross), with occasional probabilistic flips to maintain uncertainty (participants were informed that feedback was probabilistic but were not given exact probabilities). The first two blocks served as calibration (feedback probability fixed at 0.8 for correct responses). Based on each participant’s calibration accuracy, subsequent blocks used individually adjusted correct-feedback probabilities of 0.7, 0.8, or 0.9 (thresholds: *<* 0.56 ⇒ 0.9; 0.56–*<* 0.70 ⇒ 0.8; ≥0.70 ⇒ 0.7), ensuring sustained learning pressure across the session. Reward magnitude was fixed at 1.0 “point” for each correct outcome. These points were then converted into monetary payment according to performance-dependent scaling factors. Specifically, participants started with a base rate of 50 CHF, and their accumulated points were translated into an additional bonus ranging from 0 to 48 CHF. During the two initial calibration blocks, reward probabilities were fixed (0.8 for correct feedback) and a default conversion factor was used. Based on accuracy in these blocks, participants were assigned to one of three difficulty levels (easy, medium, hard) for the subsequent blocks, which determined both the probabilistic structure of feedback (0.9, 0.8, or 0.7 probability of positive feedback) and the money conversion factor (with harder conditions yielding higher monetary weightings). This ensured that overall payment opportunities were balanced across individuals while keeping the task suitably challenging.

### Statistical-learning (stimulus frequency) structure

In addition to the reinforcement structure, we embedded a statistical structure in the frequencies of the nine possible multisensory combinations (3 visual species ×3 calls/dances). Frequencies followed a 3 × 3 “joint probability” matrix with three levels: **common** (*p* = 0.50 of the trials for that visual species), **occasional** (*p* = 0.35), and **rare** (*p* = 0.15). Specifically, the three common pairs were (0, *A*), (1, *B*), and (2, *C*); the three occasional pairs were (0, *B*), (1, *C*), and (2, *A*); and the three rare pairs were (0, *C*), (1, *A*), and (2, *B*).

Trial sequences were generated algorithmically to meet three constraints: (1) within every 20-trial segment of a block, pair counts matched the expected ratios exactly (10 common, 7 occasional, 3 rare events); (2) across the full 60 trials of a block, frequencies matched the joint-probability matrix precisely; and (3) immediate repetitions in either unisensory modality were avoided (e.g., if the current trial was visual “1” with call “A,” the following trial could not share the same visual or the same auditory/tactile element).

These constraints ensured that the statistical structure was both robust and evenly distributed, while preventing trivial strategies such as detecting simple repetitions. Importantly, participants were never informed about the frequency manipulation, and the statistical structure was independent of reward contingencies, allowing statistical learning to be probed implicitly.

### Computational models

#### Reinforcement Learning

To model reinforcement learning computations that are explicitly required for task performance, we employed Q-learning with two 3 × 3 Q-value tables. Each table represented the Q-values for the two possible actions (attract or not attract) across the nine multisensory stimuli. This setup allowed us to estimate the value of each action for each stimulus pairing, as shown in Figure 1 from the main text. The green cells in the figure represent the pairings most likely to attract a partner (and therefore requiring a “yes” response; please note that rewards were (or were not) given for both correct (or incorrect) “yes” and “no” responses, so that rewards were independent of statistical structure and response type). Q-learning assumes that participants update their Q-values based on the RPE which is the difference between the expected value and the observed reward. Unless otherwise specified, unchosen values remained unchanged. In addition to signed RPEs, we also derived uRPEs, defined as the absolute magnitude of the RPE, indexing the degree of outcome surprise irrespective of valence [39, 37].

We tested different versions of Q-learning to capture the diverse strategies and biases that participants might have. These models included:

1. **Basic Q-learning** — The simplest model was the basic Q-learning algorithm where participants only updated the value for the chosen response, with one learning rate *α* for all possibilities [40, 20].

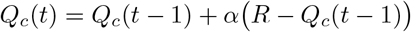

where *Q*_*c*_ denotes the value of the chosen option for the presented multisensory state in that trial. The Q-values were initialized to 0.5 for both actions in this and all other models, except for versions of the Vinit model where initial values were treated as free parameters.
2. **Asym** — Studies have suggested that people learn differently from positive and negative RPEs [41, 42, 43, 44]. To incorporate this, we designed a model with separate learning rates for positive and negative feedback:

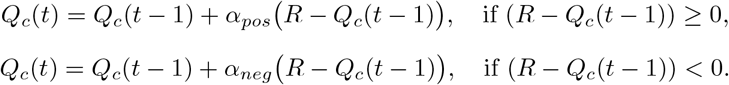
3. **Transfer** — Given the task structure, each trial conveyed information not only about the presented multisensory stimulus but also about related stimuli that shared one unisensory feature. For example, if A1 attracted a partner, then A2, A3, B1, and C1 could be inferred to be less likely. To model this inference, we allowed related, unchosen pairings to be updated with a scaled parameter:

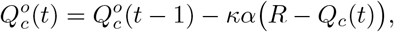

where 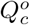 denotes the value of a related option, and *κ* ≤ 1 scales the transfer update so that related stimuli are never updated more strongly than the chosen stimulus.
4. **Vinit** — This model captured initial biases in action preferences. Instead of fixing Q-values at 0.5, the initial Q-values for both action tables were treated as free parameters, allowing us to model a priori action propensities. The free parameters were *Q*_0_, *Q*_1_, *α*. Updating followed the same rule as in basic Q-learning.
5. **TransferVinit** — This model combined the Transfer and Vinit features, i.e., transfer updates were applied and initial Q-values were free parameters.

All of the above models had parameter recovery correlations above 0.6 between simulated and recovered parameters. Therefore, their parameters were considered recoverable. We also explored additional variants, such as models with asymmetric learning rates for actions, Pearce–Hall–style adaptive learning, and dynamic learning rates, but these failed to reach the recovery threshold (*r <* 0.6) and were excluded from further fitting.

The unrecoverable models included:

- **Extra** — Asymmetric learning rates for the two different actions:

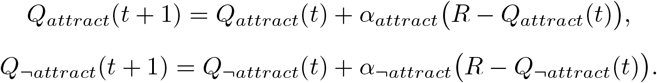
- **Pearce** — A Pearce–Hall style model [65] with trial-wise adaptive learning rates:

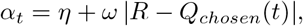

where *η* is the base learning rate and *ω* modulates sensitivity to prediction errors.
- **Dyna** — A dynamic learning-rate model:

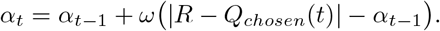

In the end, only the five models with recoverable parameters (Basic, Asym, Transfer, Vinit, TransferVinit) were fitted. All models were fit to each participant’s trial-by-trial choices via maximum likelihood (SciPy v1.10; L-BFGS-B; multiple random initializations). Model evidence was approximated with the Bayesian Information Criterion (BIC), computed from the maximum log-likelihood and number of free parameters. For model selection, we summed BIC across the six blocks per participant and chose the lowest-BIC model as that participant’s winner. Of the five recoverable models, three accounted for all participants’ data: **Basic Q-learning (n = 31), Transfer (n = 16), and Asymmetric learning rates (n = 11)**; **no participants** were best fit by **Vinit** or **Transfer+Vinit**. The per-participant winning model was then used to generate trial-wise value estimates, signed RPEs, and uRPEs for all behavioral and fMRI analyses.

### Statistical Learning

To model statistical learning, we leveraged the probabilistic structure of the task. Each visual species could be paired with one of three calls/dances, resulting in nine possible multisensory combinations. Frequencies followed a 3 × 3 joint-probability matrix with three levels: common (*p* = 0.50), occasional (*p* = 0.35), and rare (*p* = 0.15). This structure yielded a 50/50 split between pairings that were most likely versus least likely to attract a partner, preventing participants from adopting a trivial strategy of selecting the same action throughout the experiment.

We modeled statistical learning using a Bayesian updating framework without any free parameters, following [33]. The model assumes a uniform prior over the nine possible stimulus combinations at the start of each block. On every trial, beliefs about the probability of encountering each stimulus *i* are updated, resulting in a Dirichlet posterior that gradually converges toward the true empirical frequencies.

On each trial *t*, the belief for stimulus *i* is given by:

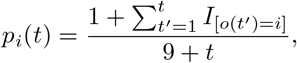

where 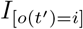 is an indicator function equal to 1 if stimulus *i* was observed on trial *t*^*′*^, and 0 otherwise. This formulation reflects Bayesian updating with a uniform prior and counts-based likelihood.

To quantify statistical learning, we computed the Shannon surprise of the observed stimulus *o* on trial *t*:

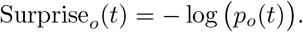

Lower prior belief *p*_*o*_(*t*) results in higher surprise. Trial-wise surprise values were then used as a proxy for statistical learning, with the hypothesis that more surprising stimuli would elicit longer response times.

### Behavioral analysis

To assess how reinforcement and statistical learning signals influenced behavior, we fit a linear mixed-effects regression model predicting trial-wise response times:

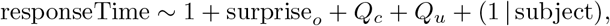

where surprise_*o*_ is the Shannon surprise of the presented stimulus on trial *t, Q*_*c*_ and *Q*_*u*_ are the chosen and unchosen action values from the fitted RL model, and (1 |subject) specifies subject-level random intercepts.

All predictors were z-scored prior to analysis to enable direct comparison of regression coefficients. Mixed-effects models were estimated in R using the <monospace>lmerTest</monospace> package, called from Python via <monospace>rpy2</monospace>. Statistical significance was evaluated using Satterthwaite’s approximation for degrees of freedom as implemented in <monospace>lmerTest</monospace>.

### MRI acquisition and preprocessing

Functional and anatomical MRI data were acquired at the Laboratory for Social and Neural Systems Research, University Hospital Zurich, using a Philips Achieva 3T whole-body MR scanner equipped with a 32-channel head coil. Functional images were collected across six runs using a T2*-weighted gradient-recalled echo-planar imaging (EPI) sequence (222 volumes + 5 dummy scans; flip angle = 90°; repetition time, TR = 2334 ms; echo time, TE = 30 ms; matrix size = 80 × 78; field of view = 240 × 240 mm; in-plane resolution = 3 mm; 40 axial slices, slice thickness = 3 mm, slice gap = 0.5 mm; SENSE acceleration factor = 1.5 in the phase-encoding direction; acquisition time = 8:54 min).

A high-resolution T1-weighted structural image was acquired using an MPRAGE sequence (field of view = 256 × 256 × 170 mm; 1 mm isotropic resolution; inversion time, TI = 2800 ms; 170 sagittal slices; flip angle = 8°; TR = 8.3 ms; TE = 3.9 ms; SENSE acceleration factor = 2 in the left–right direction; acquisition time = 5:35 min). In addition, diffusion tensor imaging (DTI) data were acquired (field of view = 224 × 224 × 118 mm; 2 mm isotropic resolution; 59 axial slices; TR = 9000 ms; 64 non-collinear diffusion-weighted directions with b = 1000 s/mm^2^, plus one non-diffusion-weighted image with b = 0 s/mm^2^; acquisition time = 10:03 min).

Preprocessing of the fMRI data was performed using *fMRIPrep* 20.2.3 [99] (RRID:SCR_016216), which is based on Nipype 1.6.1 [100] (RRID:SCR_002502) and incorporates routines from SPM. The pipeline included realignment and unwarping, slice-timing correction, coregistration, segmentation, and normalization. A full description of preprocessing steps is provided in the Appendix. Finally, spatial smoothing was applied using SPM12, with a 6 mm full-width-at-half-maximum (FWHM) Gaussian kernel to reduce noise and account for residual inter-individual differences in anatomy during group-level analyses.

### First-Level Analysis

To model task-related neural activity, we specified a general linear model (GLM) in SPM12 (build 7771) implemented in MATLAB (R2023b, version 23.2.0.2668659). The GLM was designed to capture blood-oxygen-level-dependent (BOLD) responses associated with audiovisual and visuo-tactile statistical and reinforcement learning. The design matrix included six motion parameters (three translation and three rotation estimates) to account for head motion artifacts, the first five aCompCor components derived from fMRIPrep to model noise related to physiological and non-neural signals, and physiological noise regressors (cardiac and respiratory signals) estimated using the PhysIO Toolbox for MATLAB.

The task-related regressors included the choice period, modeled as an epoch with a duration equal to the participant’s response time and parametrically modulated by Shannon surprise and the value of the chosen option. The feedback period was modeled as a fixed-duration epoch (1 second) parametrically modulated by RPE and uRPE. All parametric modulators were z-standardized (mean-centered and scaled to unit variance) to ensure comparability across participants and conditions. No orthogonalization was applied to the regressors of interest to preserve shared variance between predictors. To verify that these regressors captured distinct computational signals, we quantified their trial-wise correlations across participants. Mean Pearson correlations were very small, confirming minimal collinearity. Mean Pearson correlations were small, confirming minimal collinearity between the learning signals. Across subjects, mean coefficients of determination were RPE–SPE, *r*^2^ = 0.007±0.007; uRPE–SPE, *r*^2^ = 0.045±0.027; and RPE–uRPE, *r*^2^ = 0.110±0.118. These low shared variances indicate that the three learning signals were largely independent and suitable for simultaneous inclusion as separate regressors in the fMRI general linear model (GLM). The GLM design further accounted for any residual shared variance among regressors.

For each parametric modulator, we defined first-level contrasts for all runs vs. implicit baseline, audio runs vs. baseline, tactile runs vs. baseline, and audio runs vs. tactile runs.

### Group Analysis

For second-level inference, we employed non-parametric permutation testing using the SnPM toolbox (v13.01). This approach involved 5,000 permutations to generate a null distribution for statistical testing, cluster-wise inference with family-wise error (FWE) correction at *p <* 0.05 to control for multiple comparisons. We tested both positive and negative contrasts for RPE, Shannon surprise, and uRPE. To assess modality-specific effects, we also computed direct contrasts between audiovisual and visuotactile conditions for each variable of interest.

Cluster-forming thresholds were optimized for each analysis to balance cluster overlap and statistical sensitivity, with cluster-forming thresholds set at *t >* 8.0 for RPE, *t >* 3.1 for Shannon surprise, and *t >* 4.0 for uRPE.

### ROI Analysis

For the region-of-interest (ROI) analysis, we defined each ROI as the set of all voxels in MNI space that survived cluster-wise family-wise error (FWE) correction (at p ¡ 0.05) and exceeded the variable-specific cluster-forming threshold (see previous section). These voxels, identified in the second-level group analysis, formed the functional masks for each ROI. From each mask, we extracted the mean beta estimate per participant for the audiovisual and visuotactile regressors, focusing on the contrasts of interest: RPE, Shannon surprise, and uRPE. By comparing these mean modality-specific beta estimates across conditions, we directly tested for differences in neural activation patterns related to RPE, surprise, and uRPE between audiovisual and visuotactile learning, while accounting for multiple comparisons through cluster-based thresholding.

## Code availability

All task scripts, computational modeling code, and analysis pipelines used in this study are publicly available at: https://github.com/ruffgroup/multlearn

The repository contains the scripts required to reproduce the behavioral analyses, computational modeling, and neuroimaging results reported in this manuscript.

## Data availability

The anonymized MRI dataset supporting the findings of this study has been deposited on Open-Neuro and is publicly available at: https://openneuro.org/datasets/ds007436/versions/1.0.0

Data are organized according to the Brain Imaging Data Structure (BIDS) standard and include all raw and preprocessed imaging data required to reproduce the reported analyses.

## A fMRIPrep

The preprocessing steps for the anatomical image were as follows:

A total of 1 T1-weighted (T1w) images were found within the input BIDS dataset.The T1-weighted (T1w) image was corrected for intensity non-uniformity (INU) with N4BiasFieldCorrection (Tustison et al. 2010), distributed with ANTs 2.3.3 (Avants et al. 2008, RRID:SCR_004757), and used as T1w-reference throughout the workflow. The T1w-reference was then skull-stripped with a Nipype implementation of the antsBrainExtraction.sh workflow (from ANTs), using OA-SIS30ANTs as target template. Brain tissue segmentation of cerebrospinal fluid (CSF), white-matter (WM) and gray-matter (GM) was performed on the brain-extracted T1w using fast (FSL 5.0.9, RRID:SCR_002823, Zhang, Brady, and Smith 2001). Brain surfaces were reconstructed using recon-all (FreeSurfer 6.0.1, RRID:SCR_001847, Dale, Fischl, and Sereno 1999), and the brain mask estimated previously was refined with a custom variation of the method to reconcile ANTs-derived and FreeSurfer-derived segmentations of the cortical gray-matter of Mindboggle (RRID:SCR_002438, Klein et al. 2017). Volume-based spatial normalization to one standard space (MNI152NLin2009cAsym) was performed through nonlinear registration with antsRegistration (ANTs 2.3.3), using brain-extracted versions of both T1w reference and the T1w template. The following template was selected for spatial normalization: ICBM 152 Nonlinear Asymmetrical template version 2009c [Fonov et al. (2009), RRID:SCR_008796; TemplateFlow ID: MNI152NLin2009cAsym],

For each of the 6 BOLD runs found per subject (across all tasks and sessions), the following preprocessing was performed. First, a reference volume and its skull-stripped version were generated using a custom methodology of fMRIPrep. A B0-nonuniformity map (or fieldmap) was estimated based on a phase-difference map calculated with a dual-echo GRE (gradient-recall echo) sequence, processed with a custom workflow of SDCFlows inspired by the epidewarp.fsl script and further improvements in HCP Pipelines (Glasser et al. 2013). The fieldmap was then co-registered to the target EPI (echo-planar imaging) reference run and converted to a displacements field map (amenable to registration tools such as ANTs) with FSL’s fugue and other SDCflows tools. Based on the estimated susceptibility distortion, a corrected EPI (echo-planar imaging) reference was calculated for a more accurate co-registration with the anatomical reference. The BOLD reference was then co-registered to the T1w reference using bbregister (FreeSurfer) which implements boundary-based registration (Greve and Fischl 2009). Co-registration was configured with six degrees of freedom. Head-motion parameters with respect to the BOLD reference (transformation matrices, and six corresponding rotation and translation parameters) are estimated before any spatiotemporal filtering using mcflirt (FSL 5.0.9, Jenkinson et al. 2002). BOLD runs were slice-time corrected using 3dTshift from AFNI 20160207 (Cox and Hyde 1997, RRID:SCR_005927). The BOLD time-series were resampled onto the following surfaces (FreeSurfer reconstruction nomenclature): fsaverage, fsnative. The BOLD time-series (including slice-timing correction when applied) were resampled onto their original, native space by applying a single, composite transform to correct for head-motion and susceptibility distortions. These resampled BOLD time-series will be referred to as preprocessed BOLD in original space, or just preprocessed BOLD. The BOLD time-series were resampled into standard space, generating a preprocessed BOLD run in MNI152NLin2009cAsym space. First, a reference volume and its skull-stripped version were generated using a custom methodology of fMRIPrep. Several confounding time-series were calculated based on the preprocessed BOLD: framewise displacement (FD), DVARS and three region-wise global signals. FD was computed using two formulations following Power (absolute sum of relative motions, Power et al. (2014)) and Jenkinson (relative root mean square displacement between affines, Jenkinson et al. (2002)). FD and DVARS are calculated for each functional run, both using their implementations in Nipype (following the definitions by Power et al. 2014). The three global signals are extracted within the CSF, the WM, and the whole-brain masks. Additionally, a set of physiological regressors were extracted to allow for component-based noise correction (CompCor, Behzadi et al. 2007). Principal components are estimated after high-pass filtering the preprocessed BOLD time-series (using a discrete cosine filter with 128s cut-off) for the two CompCor variants: temporal (tCompCor) and anatomical (aCompCor). tCompCor components are then calculated from the top 2% variable voxels within the brain mask. For aCompCor, three probabilistic masks (CSF, WM and combined CSF+WM) are generated in anatomical space. The implementation differs from that of Behzadi et al. in that instead of eroding the masks by 2 pixels on BOLD space, the aCompCor masks are subtracted a mask of pixels that likely contain a volume fraction of GM. This mask is obtained by dilating a GM mask extracted from the FreeSurfer’s asegsegmentation, and it ensures components are not extracted from voxels containing a minimal fraction of GM. Finally, these masks are resampled into BOLD space and binarized by thresholding at 0.99 (as in the original implementation). Components are also calculated separately within the WM and CSF masks. For each CompCor decomposition, the k components with the largest singular values are retained, such that the retained components’ time series are sufficient to explain 50 percent of variance across the nuisance mask (CSF, WM, combined, or temporal). The remaining components are dropped from consideration. The head-motion estimates calculated in the correction step were also placed within the corresponding confounds file. The confound time series derived from head motion estimates and global signals were expanded with the inclusion of temporal derivatives and quadratic terms for each (Satterthwaite et al. 2013). Frames that exceeded a threshold of 0.5 mm FD or 1.5 standardised DVARS were annotated as motion outliers. All resamplings can be performed with a single interpolation step by composing all the pertinent transformations (i.e. head-motion transform matrices, susceptibility distortion correction when available, and co-registrations to anatomical and output spaces). Gridded (volumetric) resamplings were performed using antsApplyTransforms (ANTs), configured with Lanczos interpolation to minimize the smoothing effects of other kernels (Lanczos 1964). Non-gridded (surface) resamplings were performed using mri vol2surf (FreeSurfer).

## Primary sensory regions

### A.1 Contrast 1: RPE

**Table 1.** Clusters that correlate with RPE, cluster-forming threshold T = 8.

| Cluster Label | Voxels | MAX<br>T | MAX<br>X<br>(mm) | MAX<br>Y<br>(mm) | MAX<br>Z<br>(mm) | P<br>FWE-CORR. |
| --- | --- | --- | --- | --- | --- | --- |
| Caudate, Putamen R | 391 | 19.29 | 18 | 14 | -12 | 0.0002 |
| Caudate, Putamen L | 273 | 16.22 | -16 | 14 | -12 | 0.0002 |
| Angular Gyrus L | 66 | 11.78 | -46 | -66 | 44 | 0.0002 |
| Visual cortex (Mid, Lat Occipital) L | 139 | 11.78 | -46 | -82 | 2 | 0.0002 |
| vmPFC | 213 | 11.05 | 8 | 54 | -8 | 0.0002 |
| Visual Cortex (Lingual, Fusiform) L | 61 | 10.80 | -22 | -88 | -12 | 0.0002 |
| Cerebellum R | 111 | 10.67 | 42 | -70 | -40 | 0.0002 |

### A.2 Contrast 5: Surprise

**Table 2.** Clusters that correlate with surprise, cluster-forming threshold T = 3.1.

| Cluster Label | Voxels | MAX T | MAX X<br>(mm) | MAX Y<br>(mm) | MAX Z<br>(mm) | P FWE-CORR. |
| --- | --- | --- | --- | --- | --- | --- |
| Precuneus L | 201 | 6.01 | -4 | -54 | 40 | 0.0094 |
| DLPFC L | 103 | 4.98 | -28 | 24 | 48 | 0.0284 |
| Angular Gyrus R | 90 | 4.83 | 48 | -72 | 37 | 0.0354 |
| Angular gyrus L | 111 | 4.68 | -36 | -82 | 40 | 0.0082 |

### A.3 RL negative T-value neural signatures

**Figure S1.**
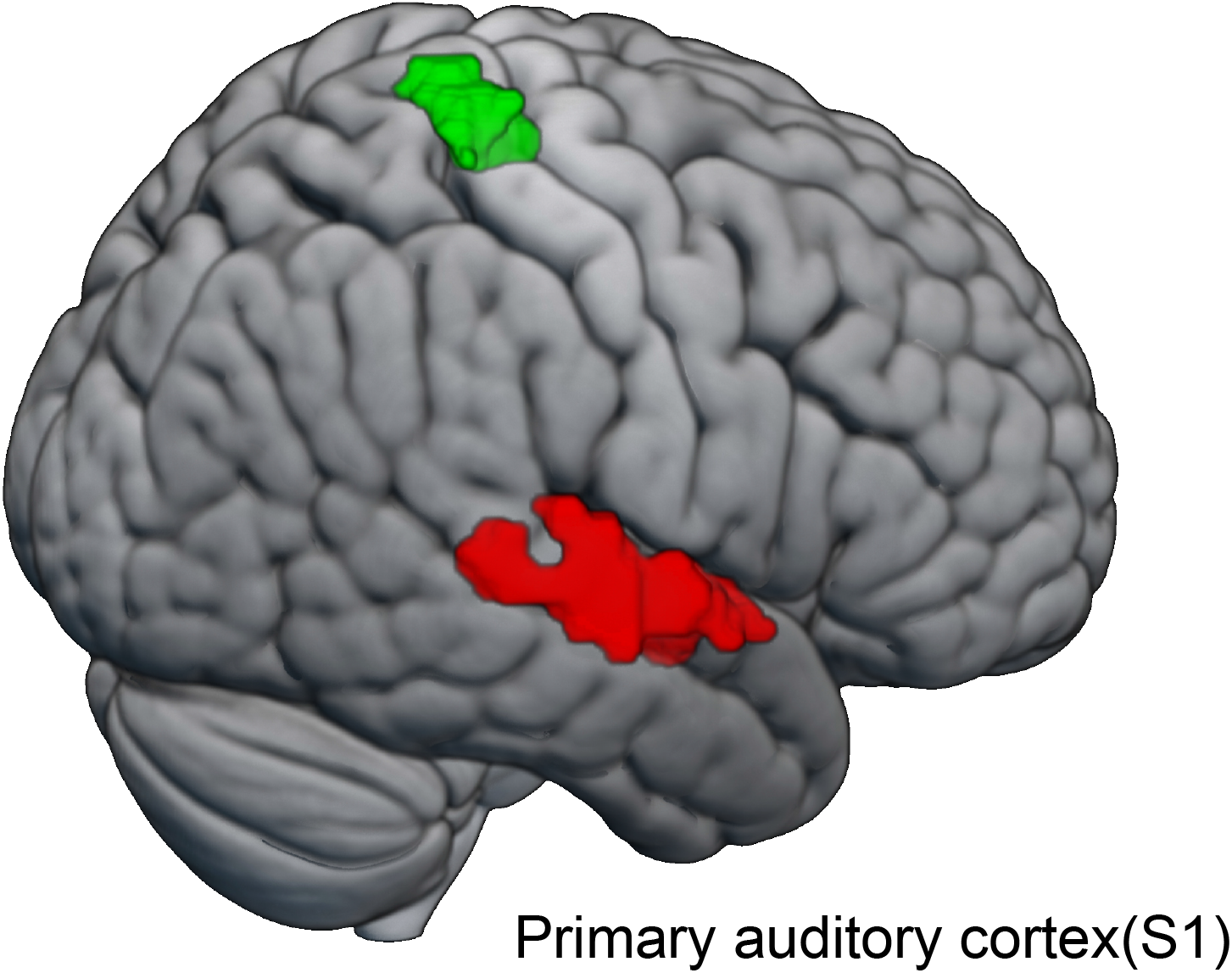
Primary sensory regions that are found with the contrast choice_audiovisual_ *>* choice_visuotactile_, thresholded at t=3.1

### A.4 Unsigned RPE

**Table 3.** Clusters that correlate with unsigned RPE (URPE), cluster-forming threshold T = 4.0.

| Cluster Label | Voxels | MAX<br>T | MAX<br>X<br>(mm) | MAX<br>Y<br>(mm) | MAX<br>Z<br>(mm) | P<br>FWE-CORR. |
| --- | --- | --- | --- | --- | --- | --- |
| dmPFC R | 142 | 8.02 | 2 | 32 | 37 | 0.0010 |
| Insula R | 116 | 7.25 | 36 | 24 | -5 | 0.0012 |
| Insula L | 31 | 5.47 | -34 | 26 | -2 | 0.0088 |
| Frontal Middle Gyrus R | 123 | 4.68 | 32 | 60 | 2 | 0.010 |
| Orbital part of the Inferior Frontal Gyrus R | 13 | 5.32 | 50 | 36 | -12 | 0.032 |
| Superior Frontal Sulcus R | 126 | 5.05 | 48 | 24 | 37 | 0.0010 |
| Inferior Parietal Lobule R | 24 | xxx | 44 | -46 | 48 | 0.0110 |

